# Unveiling the role of miRNAs from MSC-EVs in neuroinflammation and behavioral impairments induced by chronic alcohol consumption

**DOI:** 10.1101/2024.10.03.616225

**Authors:** Susana Mellado, Najoua Touahri, Sandra Montagud-Romero, Carla Perpiñá-Clérigues, Francisco García-García, Victoria Moreno-Manzano, Consuelo Guerri, Marta Rodríguez- Arias, María Pascual

## Abstract

Extracellular vesicles derived from mesenchymal stromal cells (MSC-EVs) have emerged as a promising form of regenerative and immunomodulatory therapy; indeed, micro (mi)RNAs contained within MSC-EVs modulate target gene expression and impact disease-associated pathways. Chronic alcohol consumption results in neuroinflammation, brain damage, and impaired cognition; in this study, we asked whether the repeated intravenous administration of MSC-EVs could ameliorate neuroinflammation and behavioral impairment induced by chronic alcohol consumption in mice. MSC-EVs diminished the increased binding of a micro-positron emission tomography tracer (^18^F-FDG) when analyzing whole-brain 3D images and brain coronal sections of ethanol-treated mice. MSC-EV administration protected against ethanol-induced proinflammatory gene upregulation, cognitive dysfunction, and addictive-like behavior. miRNA sequencing data from MSC-EVs helped to reveal the elevated expression of EV-derived miR-483-5p and miR-140-3p in the brains of ethanol-treated mice following MSC-EV administration. In addition, MSC-EVs modulated the expression of pro-inflammatory-related miRNA target genes (e.g., Socs3, Tnf, Mtor, Atf6) in the brains of ethanol-treated mice. These results suggest that MSC-EVs could function as a neuroprotective therapy to ameliorate the neuroinflammation, cognitive dysfunction, and addictive-like behavior associated with chronic alcohol consumption.

## INTRODUCTION

Mesenchymal stem cells (MSCs) have the capacity for self-renewal and multidirectional differentiation [1]; however, increasing evidence suggests that their therapeutic potential derives from secreted factors such as extracellular vesicles (EVs) [2]. MSC-EVs have garnered significant attention as promising therapeutic candidates for treating conditions such as cancer, autoimmune disease, neurological disorders, and inflammation [1]. Although MSC-EVs transfer numerous types of bioactive molecules (e.g., DNA, RNA, proteins, and lipids) to target cells to exert their therapeutic activity, evidence suggests that the delivery of micro(mi)RNAs induces the most substantial effects [3]; therefore, treatment with MSC-EVs or specific miRNAs will potentially exert similar anti-inflammatory and regenerative effects to MSCs themselves [4]. The use of big data-based transcriptome analysis combined with computational tools has defined many of the critical factors contained within MSC-EVs in terms of functional enrichment of miRNA expression patterns and gene analysis, which has contributed to the application of MSC-EVs for appropriate target diseases [4].

Chronic consumption of alcohol - one of the most commonly abused substances worldwide - leads to dependence-associated alterations to brain structure and function and prompts the development of behavioral, cognitive, and psychiatric disorders [5]. Although the underlying mechanisms of ethanol neurotoxicity remain incompletely understood, our findings revealed that ethanol induces a neuroinflammatory immune response, leading to the release of cytokines and chemokines that cause brain damage and behavioral dysfunction [6]. While the administration of anti-inflammatory compounds may have potential as a short-term treatment option for ethanol consumption, developing effective therapies for chronic alcohol abuse remains challenging due to the complex pathophysiology of alcohol dependence [7]. Our recent studies have demonstrated the potential of MSC-EVs to ameliorate the upregulation of inflammatory gene expression and NLRP3 inflammasome activation and recover the cognitive and memory dysfunctions induced by binge drinking in adolescent mice [8, 9].

Considering that MSC-EVs can alleviate symptoms associated with immune disorders and neurodegenerative processes [10, 11], the present study aims to evaluate whether the repeated intravenous administration of EVs from human adipose tissue-derived MSCs (MSC-EVs) can ameliorate the neuroinflammation, cognitive dysfunction, and addictive-like behavior induced by alcohol abuse. *In vivo* micro-positron emission tomography (microPET) studies demonstrated that MSC-EV administration diminished the increased specific binding of ^18^F-FDG in whole-brain 3D images and brain coronal sections of ethanol-treated mice. MSC-EVs also protected against the ethanol-induced upregulation of proinflammatory gene expression, memory/learning impairments, and addictive-like behavior. miRNA sequencing of MSC-EVs revealed the higher expression of EV-derived miR-483-5p and miR-140-3p in the brains of ethanol-treated mice after MSC-EV administration. In addition, MSC-EVs modulated pro-inflammatory-related miRNA target gene expression in the brains of ethanol-treated mice, suggesting that the miRNAs contained within MSC-EVs played a neuroprotective role in chronic alcohol-induced neuroinflammation and behavior impairments.

## MATERIALS AND METHODS

### MSC isolation and culture and MSC-EV isolation

Human adipose tissue was obtained from surplus fat tissue isolated during knee prosthesis operations performed on four patients under sterile conditions. All human samples were anonymized. The experimental procedure was previously evaluated and accepted by the Regional Ethics Committee for Clinical Research with Medicines and Health Products following the Code of Practice 2014/01. As exclusion criteria, no samples were collected from patients with a history of cancer or infectious (viral or bacterial) diseases. All human patients voluntarily signed an informed consent document to allow the use of the adipose samples.

MSCs were expanded, grown, and characterized, as previously described [12, 13]. To collect MSC-EVs, cell media was first collected and cleared of detached cells and cell fragments by centrifugation at 300 x g for 10 min. The supernatant was then centrifuged at 2000 x g for 10 min. Subsequently, apoptotic bodies and other cellular debris were pelleted by centrifuging the supernatant at 10000 x g for 30 min. EVs were then pelleted from the resulting supernatant at 100,000 x g for 1 h. The EV pellet was washed with phosphate-buffered saline (PBS) and centrifuged at 100,000 x g for 1 h. EVs were finally suspended in PBS at 20 µg/100 µL and stored at −80 °C.

### MSC-EV characterization by transmission electron microscopy and nanoparticle tracking analysis

Freshly isolated MSC-EVs were fixed with 2% paraformaldehyde and prepared as previously described [14]. Samples were analyzed using a transmission FEI Tecnai G2 Spirit electron microscope (FEI Europe, Eindhoven, the Netherlands) with a Morada digital camera (Olympus Soft Image Solutions GmbH, Münster, Germany). The absolute size range and concentration of EVs were measured using a NanoSight NS300 Malvern (NanoSight Ltd., Minton Park, UK), as previously described [14]. **Figure 1** displays a high peak in the total number of particles ranging between 100-200 nm, which includes the size range of EVs shown by transmission electron microscopy.

**Figure 1.**
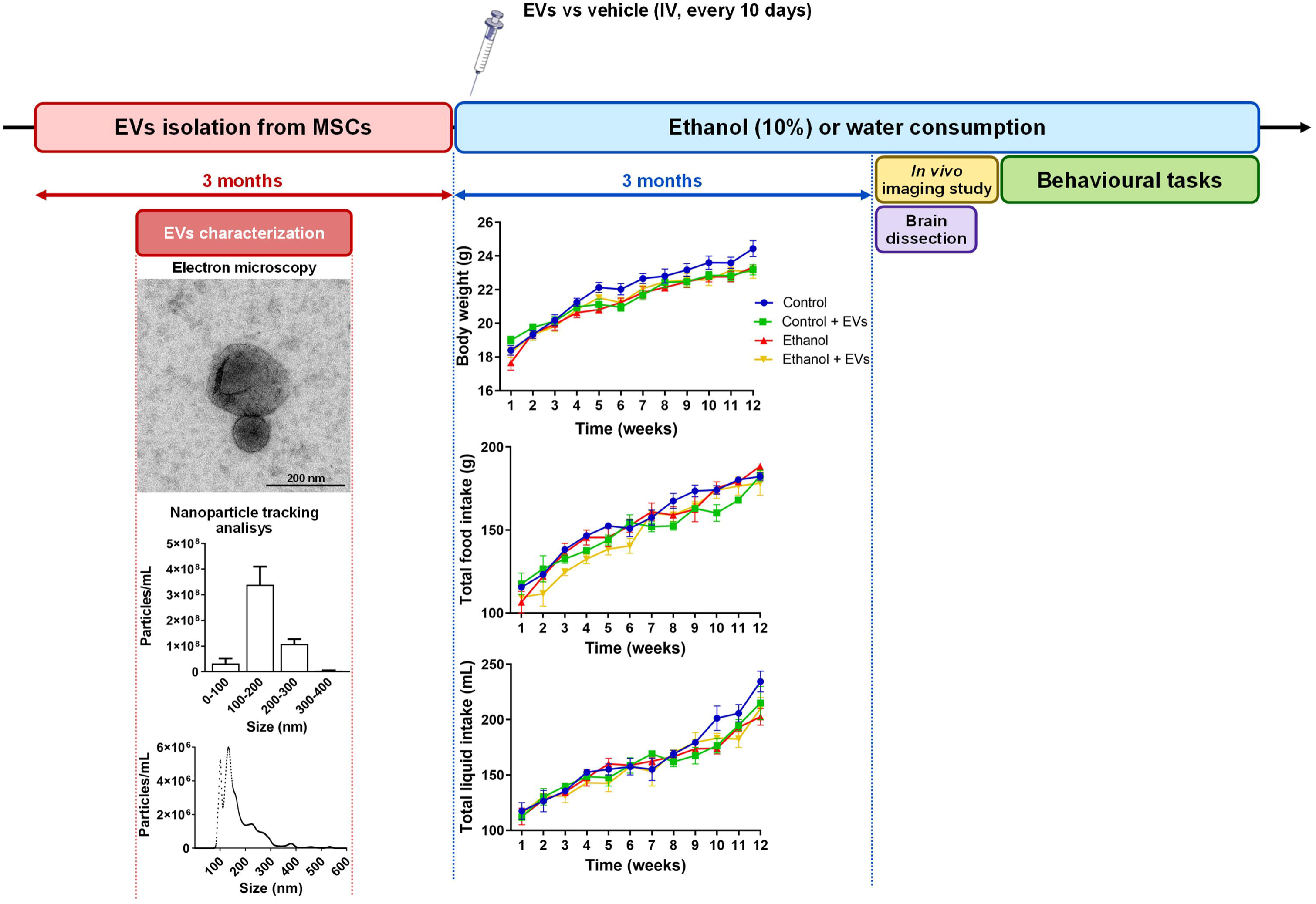
Study design. Female mice were treated with ethanol (10%) in drinking water for three months, while the control animals only drank water. MSC-EVs were injected every ten days during the three months of ethanol treatment in control and experimental mice (providing four groups: Control, Control + EVs, Ethanol, and Ethanol + EVs). MSC-EV characterization by nanoparticle tracking analysis revealed a peak between 100 and 200 nm, covering the size range of EVs revealed by transmission electron microscopy. Body weight, total liquid, and food intake were recorded during ethanol treatment. Values represent mean ± SEM, n = 12-13 mice/group. At the end of ethanol treatment, animals were sacrificed to dissect specific brain areas for gene expression analysis or used for *in vivo* imaging studies and behavioral tasksevaluations.

### Animals and treatment

Two-month-old female C57BL/6 mice (Charles River Laboratories, Wilmington, MA, USA) were used. Mice were housed (3-4 animals/cage) and maintained on a water and solid diet *ad libitum*. Environmental conditions, such as light and dark cycles (12/12 h), temperature (23 °C), and humidity (60 %), were controlled for all animals. All experimental procedures were carried out in accordance with the guidelines approved by the European Communities Council Directive (2010/63/ECC) and Spanish Royal Decree 53/2013 modified by Spanish Royal Decree 1386/2018, with the approval of the Ethical Committee of Animal Experimentation of the University of Valencia (Valencia, Spain) and the Generalitat Valenciana on January 20th, 2023 (Project identification code: 2022 VSC PEA 0294).

Mice were treated with drinking water (control) or water containing 10% (v/v) ethanol for three months. Some animals received MSC-EVs (20 µg/dose) or saline (sodium chloride, 0.9 %) via the tail vein every ten days. Animals were randomly assigned to four groups according to treatment: 1) control, 2) control + MSC-EVs, 3) ethanol, and 4) ethanol + MSC-EVs. Daily food and fluid intake in the four groups were carefully measured for three months, with no differences observed (**Figure 1**). In addition, the body weight gained during the three months remained similar in all groups (**Figure 1**), and ethanol-treated mice showed to be in optimal condition. From our previous studies using the same treatment schedules, similar blood ethanol levels were showed in the ethanol-treated groups (125 ± 20 mg/dL) [15]. After ethanol treatment, some animals were anesthetized via an intraperitoneal injection of sodium pentobarbital (60 mg/kg; Vetoquinol, Lure, France) and then sacrificed by cervical dislocation. Brains (n=8-9 mice/group) were removed, and the prefrontal cortex, striatum, and hippocampus were dissected, immediately snap-frozen in liquid nitrogen, and stored at −80 °C until used. Other animals were scanned using an in vivo fluorescent imaging system, microPET, and magnetic resonance imaging (MRI) (n=4-6 mice/group) and then used for behavioral studies (n=13-15 mice/group).

### MicroPET experiments

microPET experiments were conducted after three months of ethanol treatment in all four groups of mice. Mice were anesthetized via 3.0-4.0% isoflurane inhalation (Parsippany-Troy Hills, NJ, USA) in 100 % O_2_ and then injected intraperitoneally with 200-300□μCi of ^18^F-fluoro-2-deoxyglucose (^18^F-FDG; Advanced Accelerator Applications, Saint-Genis-Pouilly, France). Animals were returned to their housing cages, with ^18^F-FDG tracer uptake lasting approximately 30 min. After tracer uptake, animals were anesthetized via the inhalation of isoflurane mixed with 100 % O_2_ (3.0-4.0 % for anesthetic induction and 1.0-2.5 % for anesthesia maintenance). Mice were placed on a thermo-regulated bed in the prone position and scanned with the MRS*PET CLIP-ON (MR-Solutions, Guildford, UK), which contains a continuous silicon photomultiplier detector (SiPM) with a double-layer LYSO/LYSO crystal (10 mm). For radiotracer readings (45 min after ^18^F-FDG injection), 15-min list mode static acquisitions were acquired within the field of view (36.12 × 36 × 12 × 60.48 mm) centered on the mouse head. These images were obtained using the preclinical SCAN program (MR-Solutions), calibrating time, and injected dose. All data were reconstructed using the PET reconstruction program (MR-Solutions), with a field of view diameter of 34 mm, voxel size of 0.42 mm^2^, and two iterations. During radiotracer readings, the respiratory and heart rates and body temperature were monitored and kept as constant as possible. The Paxinos mouse brain atlas [16] and MRI and CT (computed tomography) templates were used to overlay the normalized images previously coregistered to the microPET image database. ^18^F-FDG binding was calculated in the whole brain and both regions (right and left) of the prefrontal cortex, striatum, nucleus accumbens, amygdala, and hippocampus using the VivoQuant 2022 (Invicro, Needham, MA, USA). Three-dimensional sets of regions of interest (ROI) were drawn using VivoQuant 2022 (Invicro) and then applied to individual ^18^F-FDG images to retrieve ^18^F-FDG uptake values in each brain. Therefore, the values of each ROI in each brain were divided by the whole-brain mean of the signal for ROI normalization. Data were expressed as ^18^F-FDG binding in kBq/cm^3^.

### Magnetic resonance imaging

Mice were deeply anesthetized as described for the microPET experiments. Imaging was carried out on MRS*DRYMAG 3.0T (MR-Solutions) and preclinical SCAN program (MR-Solution). T1-weighted sequences were used to visualize structural brain alterations in the hippocampus and cortex using MSME sequences (multi-spin multi-echo): 21.88 × 25 x mm^3^ spatial resolution. MRI images were analyzed using ImageJ software (version 1.54h, NIH, Bethesda, MD, USA).

### *In vivo* fluorescent imaging system

IVIS^®^ Lumina^™^ X5 Imaging System (Revvity, Inc., Waltham, MA, USA) was used to non-invasively visualize *in vivo* brain inflammation. The IVISense™ Pan Cathepsin 750 FAST Fluorescent Probe (ProSense^®^) (Revvity, Inc.) was used following the manufacturer’s instructions. Before the imaging session, mice received an intravenous injection of the fluorescent probe (4 nmol/mouse) dissolved in 100 µL of physiological saline. Then, mice were anesthetized with isoflurane (3.0-4.0 % for anesthetic induction and 1.0-2.5 % for anesthesia maintenance) in 100 % O_2_, placed into the imaging chamber at 37 °C, and imaged at various time points (0, 6, 24, 48 and 96 h) after probe injection. The resulting light emission was quantified using the Living Image^®^ 4.0 software (Revvity, Inc.). Data was reported as arbitrary units of radiant efficiency.

### Total RNA isolation, reverse transcription, and quantitative PCR

The frozen prefrontal cortex, striatum, and hippocampus were used for total RNA extraction. Tissues were disrupted using TRIzol (Sigma-Aldrich, St. Louis, MO, USA), and the total RNA fraction was extracted following the manufacturer’s instructions. Total mRNA and total miRNA were reverse-transcribed using the NZY First-Strand cDNA Synthesis Kit (NZYTech, Lda. Genes and Enzymes, Lisbon, Portugal) and TaqMan™ Advanced miRNA Assays (Thermo Fisher Scientific, Waltham, MA, USA).

qPCR was performed in a QuantStudio^™^ 5 Real-Time PCR System (Applied Biosystems, Waltham, MA, USA). Genes were amplified employing the AceQ^®^ qPCR SYBR Green Master Mix (NeoBiotech, Nanterre, France) following the manufacturer’s instructions. The mRNA level of the cyclophilin A housekeeping gene was used as an internal control for the normalization of analyzed genes. Specific miRNAs were amplified by the TaqMan™ Fast Advanced Master Mix (Thermo Fisher Scientific), and snRNA U6 was used as an internal control. All qPCR runs included non-template controls (NTCs). Experiments were performed in triplicate. The quantification of expression (fold change) from the Cq data was calculated by the ΔΔCq method [17] by the QuantStudio™ Design & Analysis Software (Applied Biosystems). Details of the nucleotide sequences of the used primers and miRNA assays are detailed in the Supplementary Material (**Tables S1 and S2**).

### RNA extraction from MSC-EVs, miRNA sequencing, and bioinformatics analysis

#### Total MSC-EVs RNA isolation and library preparation

Total RNA from MSC-EVs was isolated using the Total Exosome RNA Isolation Kit, following the manufacturer’s instruction (Invitrogen, Waltham, MA, USA). The construction of miRNA libraries was performed using the Small RNA-Seq Library Prep Kit for Illumina (Lexogen, Wien, Austria). Small RNA transcripts were first converted into cDNA libraries, followed by quality and quantity assessments employing High Sensitivity DNA Chips and the 2100 Bioanalyzer System (Agilent Technologies, Santa Clara, CA, USA). The final libraries were pooled in equimolar concentration for sequencing in NextSeq 550 Sequencing System (Illumina, San Diego, CA, USA) through NextSeq 500/550 v2.5 Kit (Illumina) with 1 × 150 bp read length and 400 M maximum reads per run. The raw data generated by the high-throughput sequencing of miRNA were exported as fastq files.

#### Bioinformatics/pipelines analysis

The reads were preprocessed with Cutadapt (version 4.6) [18]. After removing adapters, the trimmed sequences were aligned against the *Homo sapiens* sequences from miRBase mature.fa file [19] using Bowtie2 (version 2.5.3) [20], allowing for the detection and annotation of the mature miRNAs of interest. Following this, a count matrix of the samples was generated.

#### Functional analysis

Based on the top four most abundant miRNAs by read counts in samples that have orthologs in mice (mmu-miR-483-5p, mmu-miR-3960, mmu-miR-191-5p, mmu-miR-140-3p), the multiMiR package and database (version 1.24.0) [21] was used to obtain their validated target genes. The R version used for this analysis was 4.3.2 [22]. Protein-protein interaction networks were also constructed using the STRING web tool (https://string-db.org/). Cytoscape (v. 3.10.1) [23] was used with the Cytoscape StringApp to generate a new network using the target genes of experimentally validated miRNAs. Proteins in the network were filtered based on nervous system tissue with a threshold of 3.5, and the edge score was set at 0.6, reducing the number of target genes for subsequent functional enrichment analysis. The functional enrichment analysis was conducted using pathways from the KEGG (Kyoto Encyclopedia of Genes and Genomes), Reactome, and WikiPathways databases.

### Behavioral testing

#### Novel object recognition test

Mice performed the novel object recognition test in a black open box (24 cm × 24 cm × 15 cm) using small, non-toxic objects: two plastic boxes and a plastic toy. The task procedure is described elsewhere [8] and consists of three phases: habituation, the training session (T1), and the test session (T2). During the habituation session, mice spent 5 min exploring the open-field arena where T1 and T2 were performed. During the training session, one mouse was placed in the open-field arena containing two identical sample objects placed in the middle of the testing box for 3 min. After a 1-min retention interval, the animal was returned to the open-field arena with two objects during the test session (3 min): one object was identical to the sample, and the other was novel. The recognition index was calculated by measuring the discrimination index [D.I. = (tnovel – tfamiliar)/(tnovel + tfamiliar) × 100 %], with “t” taken as the time that each mouse spent exploring an object.

#### Passive avoidance test

The passive avoidance test employed a step-through inhibitory avoidance apparatus for mice (Ugo Basile, Comerio-Varese, Italy). This cage was made of Perspex sheets and divided into two compartments (15□×□9.5×□16.5 cm). The safe compartment was white and illuminated by a light fixture (10 W) fastened to the cage lid, whereas the “shock” compartment was dark and made of black Perspex panels. Both compartments were divided by a door that was automatically operated by sliding on the floor. The floor was made of 48 stainless steel bars (0.7 mm in diameter) placed 8 mm apart. Passive avoidance tests were carried out following the procedure previously described [8].

#### Conditioned place preference

Eight identical Plexiglas boxes with two compartments of equal size (30.7 × 31.5 × 34.5 cm high) separated by a gray central area (13.8 × 31.5 × 34.5 cm high) were employed for place conditioning. The compartments had different colored walls (black vs. white) and distinct floor textures (fine grid in the black compartment vs. wide grid in the white compartment). Four infrared light beams in each compartment of the box and six in the central area allowed the position of the mice and their crossings from one compartment to the other to be recorded. The equipment was controlled by three computers using MONPRE 2Z software (Cibertec, Madrid, Spain). Place conditioning, consisting of three phases, was carried out during the dark cycle following an unbiased procedure regarding initial spontaneous preference [24].

During the first phase (pre-conditioning - Pre-C), mice were allowed access to both apparatus compartments for 900 s per day on three consecutive days. On day three, the time spent in each compartment was recorded. Mice showing a strong unconditioned aversion (<33 % of session time; i.e., 250 s) or preference (>67 % of the session time; i.e., 650 s) for any compartment were discarded from the rest of the study. No significant differences between the time spent in the drug-paired and vehicle-paired compartments during the Pre-C phase were found.

In the second phase (conditioning), which lasted four days, mice were conditioned with 1.5 mg/kg cocaine hydrochloride (Alcaliber laboratory, Madrid, Spain) or physiological saline. The cocaine dose was selected based on previous conditioned place preference studies showing that doses below 3 mg/kg are subthreshold [25–27]. During this phase, half of the mice in each group received the drug or vehicle in one compartment, while the other half received it in the other compartment. An injection of physiological saline was administered before confining mice to the vehicle-paired compartment for 30 min. After an interval of 4 h, the mice received cocaine immediately before confinement in the drug-paired compartment for a further 30 min. The central area was made inaccessible by guillotine doors during conditioning. The dose of cocaine used during the conditioning phase was a subthreshold dose (1.5 mg/kg, proven to be ineffective in controls) to evaluate increased sensitivity to the conditioned rewarding effects of cocaine.

In the third phase (postconditioning - Post-C), which took place on day eight, the guillotine doors separating the two compartments were removed, and the time spent in each compartment by the untreated mice during a 900 s observation period was recorded. The difference in seconds between the time spent in the drug-paired compartment during the Post-C test and the Pre-C phase measures the degree of conditioning induced by the drug. A positive difference suggests that the drug has induced a preference for the drug-paired compartment, while a negative difference indicates the development of an aversion.

For extinction, mice were placed in the conditioned place preference apparatus daily, and the time spent in each compartment was measured to determine if cocaine-induced preference had disappeared. Although the group’s mean determined the day extinction was considered to have been achieved, preference was considered extinguished when a mouse spent 378 s or less in the drug-paired compartment on two consecutive days. This time was chosen based on the values of all the Pre-C tests performed in the study (mean □=□ 360s). When the preference was not extinguished in an animal, the number of days required for extinction for the group as a whole was assigned. Finally, mice were challenged with a cocaine injection once 24 h after reaching the extinction criterion, followed by a place preference test (reinstatement test).

### Statistical analysis

The results are reported as mean ± SEM. All statistical data analysis was performed using SPSS Statistics v28 (IBM Corp, Armonk, NY, USA). The Shapiro-Wilk test was used to analyze data distribution normality. A one-way ANOVA test was used as a parametric test, followed by Bonferroni’s *post* hoc test. The Mann–Whitney U test was used as a non-parametric alternative. Values of p < 0.05 were considered statistically significant.

## RESULTS

### MicroPET imaging reveals that MSC-EVs ameliorate ethanol-induced neuroinflammation

Our previous results demonstrated the therapeutic role of MSC-EVs in ameliorating the neuroinflammation induced by binge ethanol drinking in adolescent mice [8, 9]. To analyze the potential protective effects of MSC-EVs induced by chronic alcohol consumption, we first analyzed whether MSC-EVs diminished the neuroinflammation induced in our experimental model of chronic ethanol exposure using *in vivo* microPET imaging. The glucose analog ^18^F-FDG identifies localized metabolic alterations and represents a well-established imaging tracer in the differential diagnosis of neuroinflammatory processes [28] and neurodegenerative diseases [29]. **Figure 2A** shows an acquired microPET-mediated 3D image set depicting whole mouse brains in the different experimental groups. The averaged images of ^18^F-FDG uptake revealed significantly increased tracer binding in ethanol-treated mice compared to control; however, MSC-EV administration to ethanol-treated animals diminished the increased uptake of ^18^F-FDG induced by the chronic ethanol consumption to a level similar to the control (**Figure 2A**, right).

**Figure 2.**
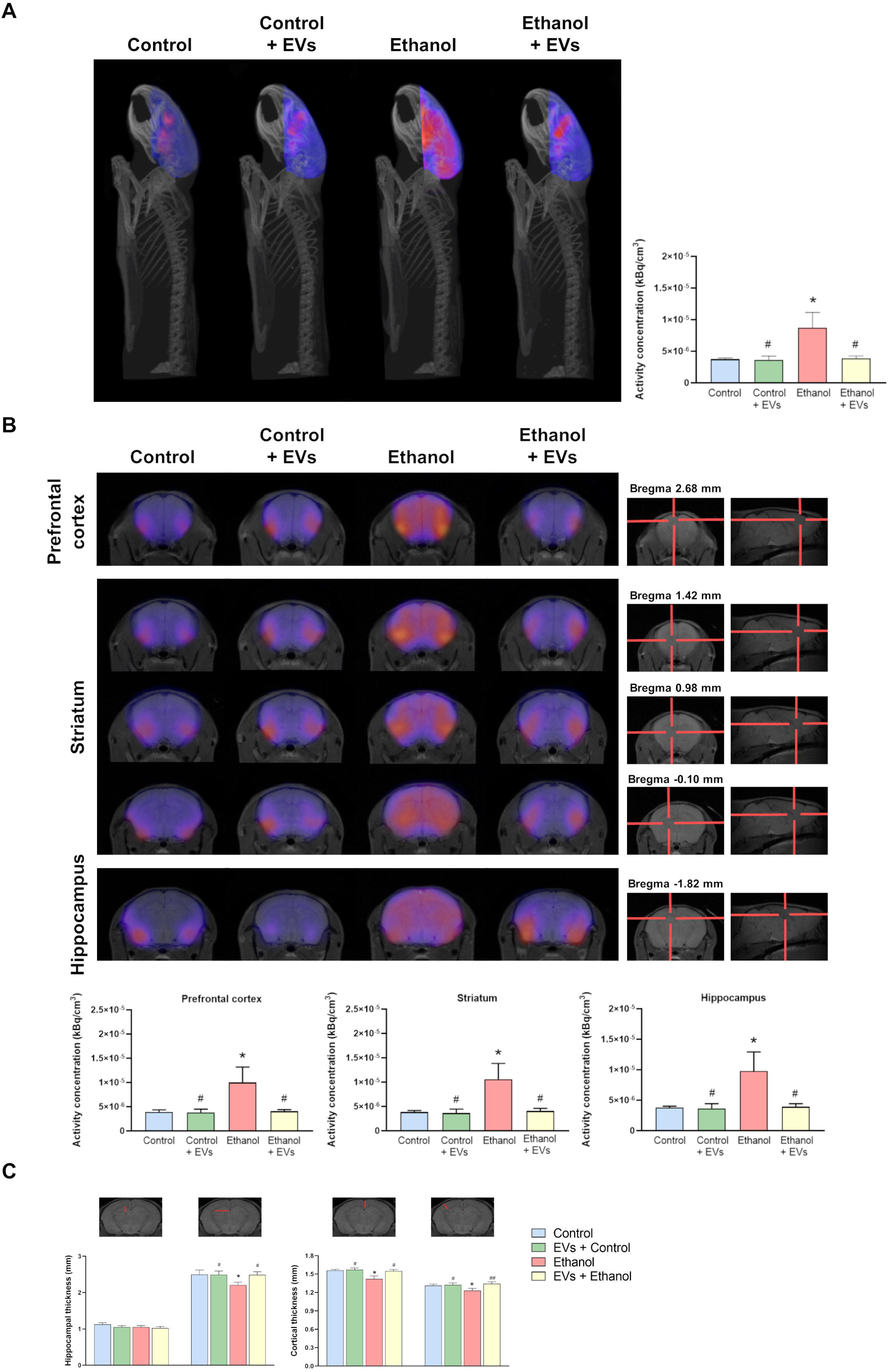
MicroPET analysis reveals that MSC-EVs decrease ethanol-induced brain ^18^F-FDG uptake in mice. **A)** Representative three-dimensional images (left) and quantification of ^18^F-FDG uptake (right) in the brains of mice treated with ethanol and/or MSC-EVs for three months (and controls). **B)** Representative coronal images (top, left) and quantification of ^18^F-FDG uptake (bottom) in the prefrontal cortex, striatum, and hippocampus of mice treated with ethanol and/or MSC-EVs for three months (and controls). Coronal and sagittal MRI templates used for coregistration in the prefrontal cortex, striatum, and hippocampus (top, right). Data represent mean ± SEM, n=4-6 mice/group. * p < 0.05 compared to control mice; # p < 0.05 and ## p < 0.01 compared to ethanol-treated mice. **C)** Representative coronal MRI images and quantification of hippocampal and cortical thickness of mice treated with ethanol and/or MSC-EVs for three months (and controls). Data represent mean ± SEM, n=4-6 mice/group. * p < 0.05 compared to their respective control group; ^#^ p < 0.05 and ^##^ p < 0.01, compared to their respective ethanol-treated group.

Further microPET analysis mapped the ^18^F-FDG distribution pattern in distinct brain areas - the prefrontal cortex, striatum, and hippocampus, which function in cognition, risk/reward, motivation, and addictive-like behavior [30–33]. We observed significantly higher specific binding of ^18^F-FDG in these brain areas when analyzing coronal brain sections of ethanol-treated mice compared to control (**Figure 2B**); furthermore, we also observed significantly decreased tracer binding in ethanol-treated mice administered with MSC-EVs compared with ethanol-treated mice to a level similar to control. We also evaluated structural hippocampal and cortical alterations using MRI (**Figure 2C**; top - representative images), as the acquired images defined the anatomic boundaries of these brain regions well [34]. While ethanol treatment significantly reduced hippocampal and cortical thickness compared to control, MSC-EV administration in ethanol-treated mice restored any significant ethanol-induced structural alterations (**Figure 2C**; bottom - quantification).

Of note, we failed to observe any significant differences between control and MSC-EV-treated animals in the microPET and MRI data (**Figure 2A-C**).

### MSC-EVs reduce inflammatory gene expression in the prefrontal cortex, striatum, and hippocampus of ethanol-treated mice

To corroborate the observed neuroprotective role of MSC-EVs in ethanol-treated mice, we next evaluated inflammation-associated gene expression in the prefrontal cortex, striatum, and hippocampus by measuring levels of Il1b, Il6, Ccl2, Ccl3, Cx3cl1, Nos2, and Cox2. **Figure 3A** demonstrates that ethanol treatment significantly upregulated Il1b, Il6, Ccl2, Ccl3, and Nos2 expression in the prefrontal cortex, striatum, and hippocampus compared to control. We observed no significant differences in the expression of these genes between MSC-EV-treated and control animals in any brain region; however, MSC-EV administration attenuated the ethanol-induced increase in inflammation-associated gene expression compared to ethanol-treated mice in the prefrontal cortex, striatum, and hippocampus.

**Figure 3.**
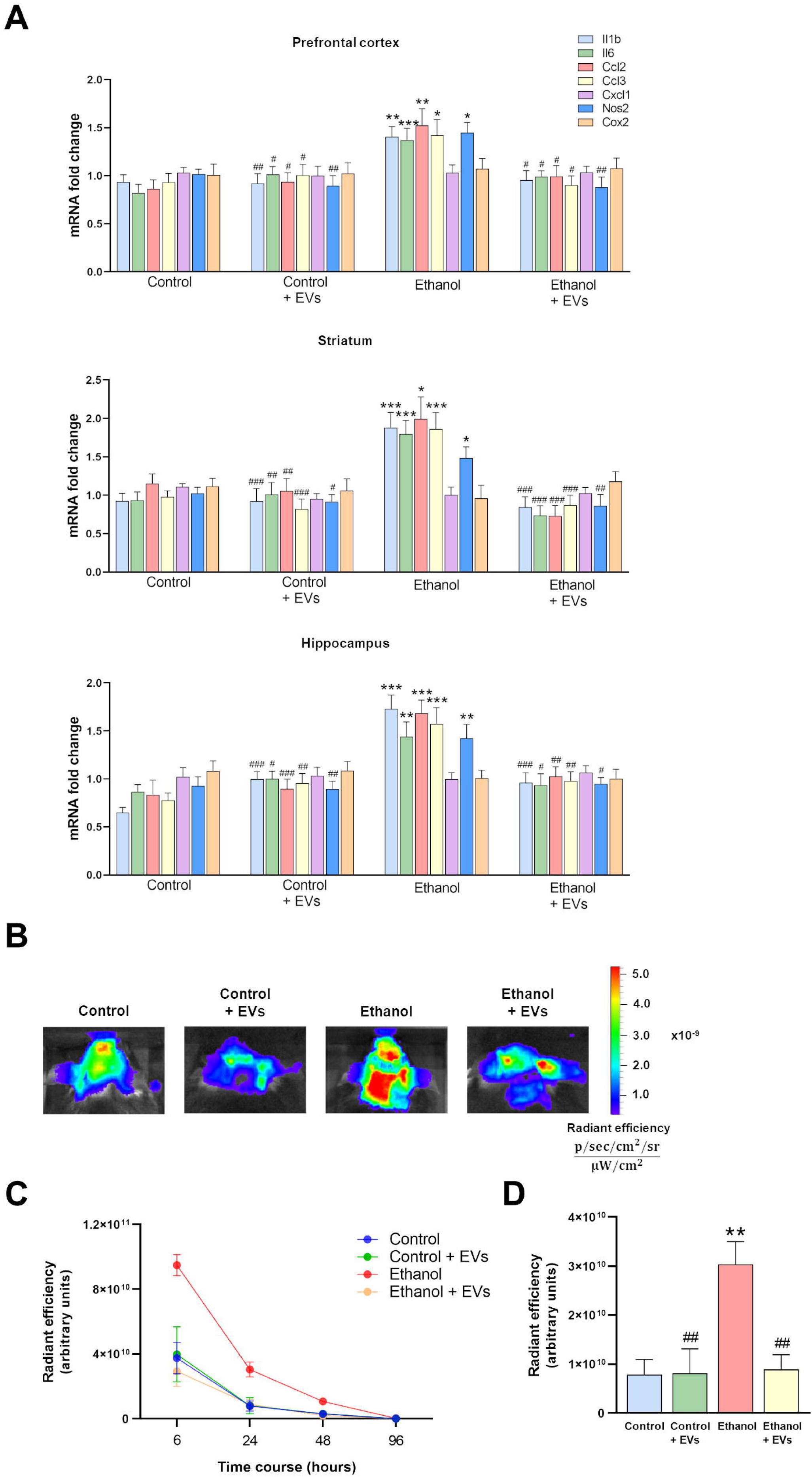
MSC-EVs decrease ethanol-induced inflammatory gene expression and *in vivo* neuroinflammation in mice. **A)** mRNA levels of Il1b, Il6, Ccl2, Ccl3, Cx3cl1, Nos, and Cox2 analyzed in the prefrontal cortex, striatum, and hippocampus of mice treated with ethanol and/or MSC-EVs for three months (and controls). Data represent mean ± SEM, n = 8-9 mice/group. * p < 0.05, ** p < 0.01, and *** p < 0.001 compared to control mice; ^#^ p < 0.05, ^##^ p < 0.01, and ^###^ p < 0.001 compared to ethanol-treated mice. **B)** Representative *in vivo* fluorescent imaging of neuroinflammation in mice treated with ethanol and/or MSC-EVs for three months (and controls). **C)** Time-course displays the fluorescent intensity at 6-, 24-, 48-, and 96-h post-injection. **D)** Fluorescence quantification at 24-h after injection of the fluorescent probe. Data represent mean ± SEM, n=4-6 mice/group. ** p < 0.01 compared to control mice; ^##^ p < 0.01 compared to ethanol-treated mice.

We then employed a non-invasive *in vivo* imaging system to monitor neuroinflammation using a fluorescent probe that measures lysosomal cathepsin activity, which correlates with inflammatory-related processes [35]. **Figure 3B** depicts representative examples of brain imaging of the fluorescent probe and the quantification at 6-, 24- and 48-h post-injection (**Figure 3C**). The absence of a fluorescent signal at 96 h post-injection demonstrated probe elimination from living mice. The fluorescent intensity significantly increased at 24-h post-injection in ethanol-treated mice compared to control (**Figure 3D**). Interestingly, MSC-EV administration at 24 h diminished the fluorescent levels in ethanol-treated animals.

### MSC-EVs restore ethanol-induced behavioral impairments

Next, we assessed whether the MSC-EV-induced amelioration of ethanol-induced neuroinflammation sufficed to restore cognitive dysfunction and addictive-like behavior in mice by performing several behavioral tasks, such as novel object recognition, passive avoidance, and conditioned place preference.

A representative trajectory of the novel object recognition test depicted in **Figure 4A** demonstrates that all groups displayed analogous trajectories while exploring both objects in the training session, spending similar time periods around each familiar object (blue squares). The test session revealed that control, MSC-EV-treated, and ethanol + MSC-EV-treated mice spent more time exploring the novel object (pink triangle) than the familiar object when compared to ethanol-treated mice, defined with higher trajectories close to the novel object than the familiar object (**Figure 4B**). We then assessed novelty recognition by calculating the discrimination index, resulting in a significant reduction in the index value in ethanol-treated mice compared to all other groups (**Figure 4C**).

**Figure 4.**
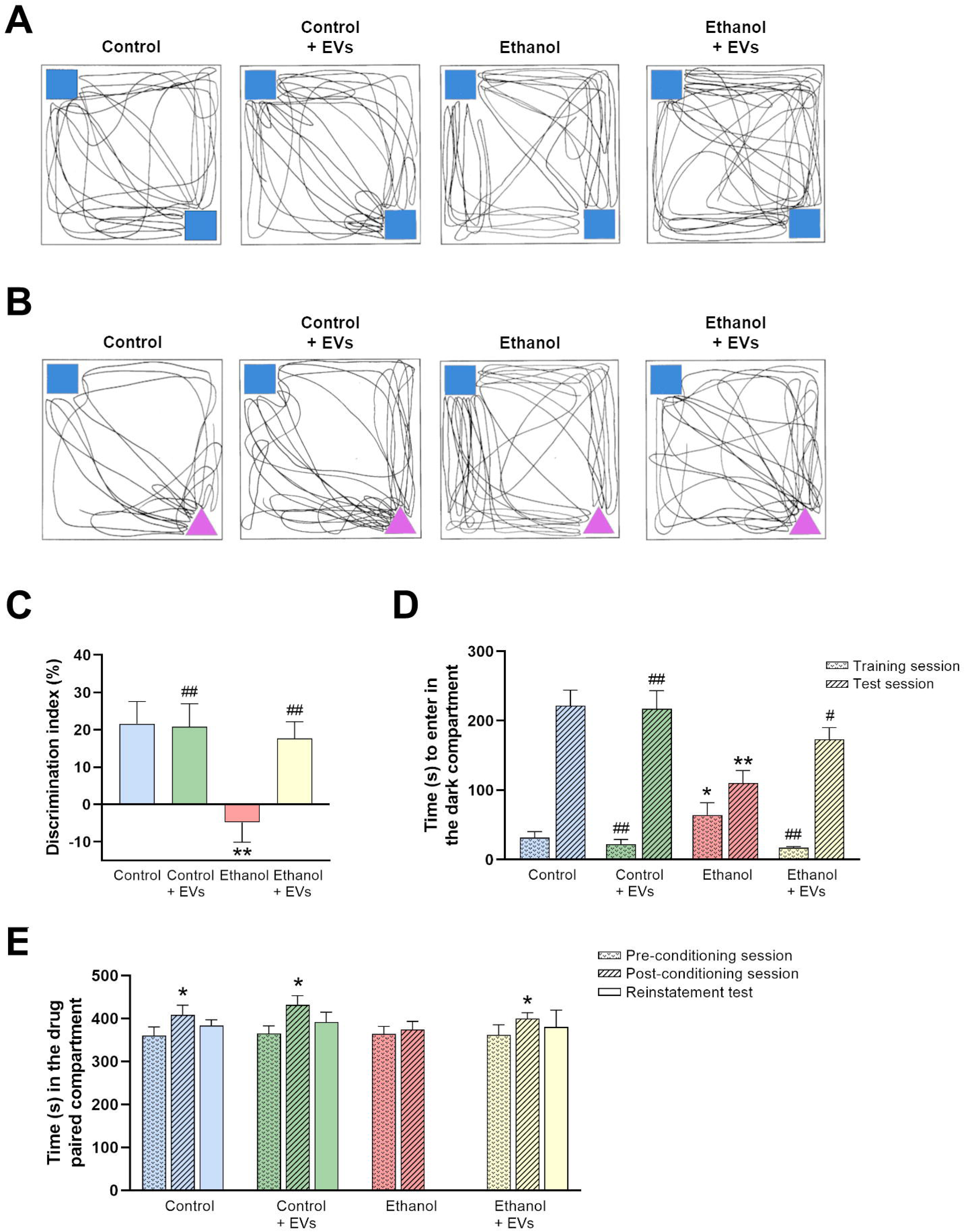
MSC-EVs restore ethanol-induced behavioral impairments in mice. A and. **B)** Representative trajectories of a single mouse from each experimental group of mice treated with ethanol and/or MSC-EVs for three months (and controls) in an open box equipped with **A**) two familiar objects (blue squares) and **B**) familiar (blue square) and novel (pink triangle) objects. **C-E)** Bar graphs represent the **C**) discrimination index during the novel object recognition task, **D**) time taken to enter the dark compartment of the passive avoidance test during the training and test session (24 h after training), and **E**) time spent in the drug-paired compartment before the conditioning sessions (Pre-C), after the conditioning sessions (Post-C), and in the reinstatement test induced by a cocaine dose of 1.5 mg/kg. Data presented as mean ± SEM, n=13-15 mice/group. * p < 0.05 and ** p < 0.01 compared to control mice; ^#^ p < 0.05 and ^##^ p < compared to ethanol-treated mice.

In the passive avoidance test (**Figure 4D**), we observed that ethanol-treated mice took longer to enter the dark compartment than mice from other groups in the training session. In addition, mice from all groups displayed longer step-through latency to enter the dark compartment during the 24-h test than the training day; however, ethanol-treated mice displayed significantly shorter latency than the other experimental groups (**Figure 4D**).

**Figure 4E** reports the time spent in the drug-paired compartment in the conditioned place preference test after a low cocaine dose of 1.5 mg/kg. The analysis revealed a significant increase in the time spent in the drug-paired compartment during the post-conditioned test compared to the pre-conditioned test in control, MSC-EV-treated, and ethanol + MSC-EV-treated mice; however, we found no significant effect in ethanol-treated mice, suggesting that these animals display less sensitivity to the low cocaine dose used in the conditioned place preference.

### miRNA sequencing revealed potential key targets of MSC-EVs

To explore the possible therapeutic mechanism of MSC-EVs regarding the amelioration of ethanol-induced neuroinflammation and behavioral impairments, we used high-throughput sequencing to identify miRNA species and their abundance in MSC-EVs. According to the sequencing data, miRNAs comprised the second-largest proportion of total RNA biotypes in MSC-EVs (Figure 5A). Figure 5B reveals the top 20 mature miRNAs in terms of abundance, where miR-483-5p, miR-3960, miR-191-5p, miR-7704, miR-7847-3p, and miR-140-3p constitute nearly 87% of total miRNA contents in MSC-EVs (Figure 5C).

**Figure 5.**
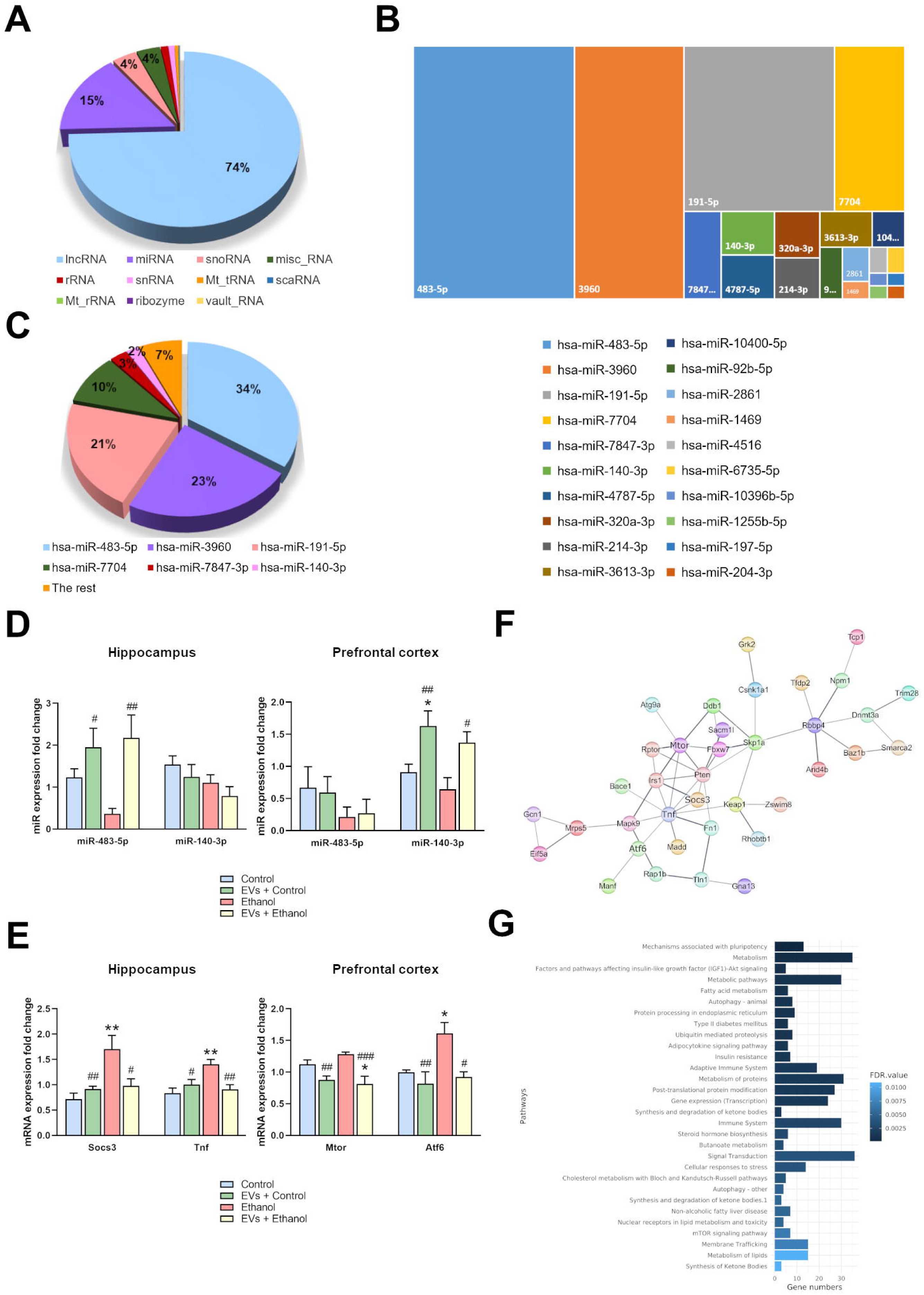
miRNA sequencing reveals potential key targets in MSC-EVs. **A)** Abundance of different RNA biotypes in MSC-EVs. **B)** The top twenty miRNAs in MSC-EVs. **C)** Percentage of the abundance of the top six miRNAs in MSC-EVs. **D)** The levels of miR-483-5p and miR-140-3p in the prefrontal cortex and hippocampus of mice treated with ethanol and/or MSC-EVs for three months (and controls). Data represents mean ± SEM, n=4-6 mice/group. * p < 0.05 compared to control mice; ^#^ p < 0.05 and ^##^ p < 0.01 compared to ethanol-treated mice. **E)** Expression levels of miR-483-5p target genes in the hippocampus (Socs3 and Tnf) and miR-140-3p target genes in the prefrontal cortex (Mtor and Atf6) of mice treated with ethanol and/or MSC-EVs for three months (and controls). Data represents mean ± SEM, n=4-6 mice/group. * p < 0.05 and ** p < 0.01 compared to control mice; ^#^ p < 0.05, ^##^ p < 0.01 and ^###^ p < 0.001 compared to ethanol-treated mice. **F)** Protein-protein interaction network for the predicted miR-483-5p and miR-140-3p target genes, filtered based on nervous system tissue, with increased edge score. Validated target genes are highlighted in bold (Mtor, Atf6, Socs3, Tnf). Line thickness indicates the strength of data support. **G)** Bar plot of the top 30 most significant pathways enriched in the genes from the previous protein-protein interaction network. The bar size indicates the number of genes involved in the pathway, and the color indicates the FDR value.

After identifying the orthologous mouse miRNAs from the six most abundant human miRNAs, we first evaluated the involvement of miR-483-5p and miR-140-3p in ameliorating inflammatory processes [36–39]. Then, we analyzed miR-483-5p and miR-140-3p expression in the prefrontal cortex and hippocampus of our experimental groups. Interestingly, we observed that the administration of MSC-EVs significantly increased the levels of both miRNAs in ethanol-treated mice in prefrontal cortex (miR-140-3p) and hippocampus (miR-483-5p), compared to ethanol-treated mice (Figure 5D).

To evaluate whether these miRNAs may mediate the neuroprotective abilities of MSC-EVs, we performed qPCR to determine the expression of some inflammatory target genes of miR-483-5p (e.g., Socs3 and Tnf) [38, 39] and miR-140-3p (e.g., Mtor, Atf6) [36, 37], obtained through a functional protein association network (**Figure S1**). Ethanol treatment significantly upregulated the expression of Mtor and Atf6 in the prefrontal cortex and Socs3 and Tnf in the hippocampus compared to the control (Figure 5E); furthermore, the administration of MSC-EVs to ethanol-treated mice significantly decreased the expression of these target genes compared to ethanol-treated mice, implying that the neuroprotective effects of MSC-EVs could be mediated by miR-483-5p and miR-140-3p and their target genes. Then, we used Cytoscape tool to analyze the target genes filtered based on the nervous system tissue, which are cooperatively affected by both miRNAs and includes Socs3, Tnf, Mtor and Atf6 (Figure 5F). The functional enrichment revealed signaling pathways, such as autophagy, cellular response to stress and immune system (Figure 5G), including TLR pathways, MyD88 cascade, Mapk signaling pathway, among others (see heatmap **Figure S2**).

## DISCUSSION

Various studies have highlighted the regenerative potential of MSC-EVs when battling neuroinflammation, brain damage, and neurodegenerative processes [10, 11]. We now provide evidence for the neuroprotective role of MSC-EVs in mouse neuroinflammation with regard to the amelioration of ethanol-induced alterations to *in vivo* microPET imaging, proinflammatory gene expression, and behavior, which could involve the transfer of miRNAs resident in MSC-EVs. Taken together, these results confirm our previous findings associated with the therapeutic function of the MSC-EVs in a mouse experimental model of binge ethanol drinking in the adolescence [8, 9].

Alcohol abuse prompts brain damage and impaired cognitive functioning, with human and animal studies supporting the role of the neuroimmune system in the effects of ethanol on the central nervous system [40, 41]. Our previous studies demonstrated that by activating the innate immune receptor TLR4 (Toll-like receptor 4) in glial cells, ethanol prompts cytokine and chemokine release and causes neuroinflammation and neural damage [42, 43]. In this study, we demonstrated that chronic ethanol consumption upregulated inflammatory gene expression (e.g., Il1b, Il6, Ccl2, Ccl3, and Nos2), which correlates with increased ^18^F-FDG uptake in the prefrontal cortex, striatum, and hippocampus. The ^18^F-FDG radiotracer has been utilized in diagnosing and treating various disorders, such as inflammatory [28] and neuroinflammatory diseases, providing information about neuronal and glial inflammation [28]. For instance, studies reported enhanced ^18^F-FDG microPET signals in mouse models of Alzheimer’s disease [44, 45] and murine neuroinflammatory conditions such as amyloidosis [46].

Part of the regenerative benefit from stem cell therapy may arise through secreted EVs, creating huge interest in applying them in clinical applications. EVs from various cell sources - including MSCs – display efficacy in models of neurological diseases, such as Alzheimer’s disease, Parkinson’s disease, or multiple sclerosis [47]. In these models, EVs attenuate proinflammatory signaling brain damage and reduce cognitive and behavioral deficits [47]. In agreement with these studies, we demonstrated that intravenous MSC-EVs administration prevents the ethanol-induced upregulation of the brain radiotracer signal in microPET and neuroinflammatory gene expression in adult mice. In addition, the therapeutic effects of MSC-EVs on neuroinflammatory responses could correlate with the restoration of spatiotemporal memory dysfunction and recognition memory deficits, as evaluated by the novel object recognition and passive avoidance tasks. Similarly, administering neural stem cell-derived EVs can alleviate lipopolysaccharide-induced chronic neuroinflammation and cognitive impairments [48]. Other studies have also demonstrated that MSC-EVs inhibit the chronic activation of NLRP3 inflammasome signaling and prevent long-term cognitive dysfunction after traumatic brain injury [49]. In Alzheimer’s disease [50] and status epilepticus [51], MSC-EVs can also improve cognitive and behavioral impairments by regulating hippocampal inflammation.

While the therapeutic role of MSC-EVs is well-known in the realm of cognitive dysfunction, their involvement in addictive processes remains unestablished. Interestingly, our results demonstrated that miRNAs encapsulated within MSC-EVs (and their target genes) protected against ethanol-induced addictive processes. Ethanol-treated mice display a lower sensitivity to a low dose of cocaine than the remaining experimental groups. Ethanol-treated mice needed a higher dose of cocaine to reproduce similar effects than the animals treated with ethanol + MSC-EVs, suggesting that the miRNAs contained in the MSC-EVs participated in the reward system. Likewise, some studies demonstrated that EV miRNAs exert protective effects against methamphetamine dependence [52], and the use of lentiviral vector-miRNA silencers abolished cocaine-induced conditioned place preference [53]. Indeed, a recent study reported that implicit cognition plays a vital role in addiction development and maintenance [54]. These cognitive processes toward addiction-related stimuli develop based on conditioning processes and are linked to cue reactivity and craving [54]. In this sense, MSC-EVs could participate in mechanisms such as neuroinflammation and myelin and synaptic alterations to repair memory/learning impairments [8], which may also participate in restoring the reinforcing properties of abused drugs.

The conditioned place preference paradigm assesses the conditioned rewarding effects of addictive substances by pairing drug effects with previously neutral cues, such as a specific apparatus compartment. Thus, contextual stimuli can acquire secondary appetitive properties when associated with cocaine, indicating abuse potential [55]. Prolonged ethanol use downregulates the reward system, reducing dopamine release and receptor sensitivity [56]. The dopaminergic system remains crucial for cocaine’s psychomotor stimulant and reinforcing effects, demonstrated through intravenous self-administration and conditioned place preference studies [57, 58]. Consequently, we hypothesize that ethanol-treated mice fail to associate a low cocaine dose with a specific compartment due to reward system downregulation, while MSC-EVs may reverse said ethanol-induced alterations, restoring cocaine-induced preference.

The evidence that MSC-EV miRNAs play a role in overall therapeutic effects is compelling [4]. Using a combination of miRNA transcriptomics and functional bioinformatics, possible neuroprotective implications of EV-miRNAs may be addressed to a target disease [4]. We employed miRNA sequencing to identify the cargoes carried by MSC-EVs and found that miR-483-5p in the hippocampus and miR-140-3p in the prefrontal cortex displayed higher expression in ethanol + MSC-EV-treated mice than ethanol-treated mice. Both miRNAs are included in the top six of all miRNAs by proportion, constituting 87 % of all mature miRNAs. Previous research revealed that exosomal miR-483-5p derived from adipose stem cells exhibited anti-inflammatory properties through the modulation of NLRP3 inflammasome in deep vein thrombosis [59]. A related study demonstrated that the MSC-derived exosome-mediated delivery of miR-140-3p ameliorated hippocampal inflammatory responses, pyroptosis, and cognitive impairment in mice with sepsis-associated encephalopathy [60]. Our results demonstrate that MSC-EVs decreased the expression of target inflammation-associated genes (Socs3, Tnf, Mtor, and Atf6) [36–39] in ethanol-treated mice, suggesting their involvement in enhancing the immune regulatory potential to fight ethanol-induced neuroinflammation.

We note certain limitations with the design/methodology of our study. Aside from difficulties obtaining the number of MSC-EVs needed, the chronic treatment and the high amount of animals required represent the primary concerns. Future efforts must clarify the roles of other functional miRNAs within MSCs-EVs and their targets in the effects of ethanol consumption. Finally, exploring additional regulatory molecular networks that participate in the impact of MSC-EVs associated with addictive-like behavior may prove fruitful.

Taken together, these novel results support the protective role of MSC-EVs in the neuroinflammatory immune response, cognitive dysfunction, and addictive-like behavior induced by chronic alcohol consumption. Identifying specific miRNAs and their target genes related to the immune response could provide a better understanding of the mechanisms involved in the neurotoxicity of alcohol abuse.

## ACKNOWLEDGMENTS

The authors would like to thank M.D. Reguilón and T. Aledón for their help in the behavioral studies. The authors also thank the Electron Microscopy Service at the Príncipe Felipe Research Centre and Stuart P. Atkinson for reviewing the manuscript.

## AUTHOR CONTRIBUTIONS

MP, SM, and MRA conceived and designed the experiments. SM, NT, SMR, and CPC performed the experiments and analyzed the data. SM, NT, SMR, CPC, and MP wrote the manuscript. SM, FGG, VMM, MRA, CG, and MP revised the manuscript. All authors read and approved the final manuscript.

